# Inclusion of processed cell metadata improves single cell sequencing analysis reproducibility and accessibility

**DOI:** 10.1101/2020.11.20.391920

**Authors:** Sidhant Puntambekar, Jay R. Hesselberth, Kent A. Riemondy, Rui Fu

## Abstract

Single cell RNA sequencing provides an unprecedented view of cellular diversity of biological systems. Thousands of scRNA-seq datasets have been generated, providing a wealth of biological data on the diversity of cell types across different organisms, developmental stages, and disease states. But while a tremendous number of publications and datasets have been generated using this technology, we found that a minority (< 25%) of studies provide sufficient information to enable direct reuse of their data for further studies. This problem is common across journals, data repositories, and publication dates. The lack of appropriate information not only hinders exploration and knowledge transfer of reported data, but also makes reproducing the original study prohibitively difficult and/or time-consuming. Correcting this problem is not easy but we encourage investigators, reviewers, journals, and data repositories to take steps to improve their standards and ensure proper documentation of these valuable datasets.

## Introduction

Single cell sequencing has empowered discoveries of cell heterogeneity and state transitions at unprecedented resolution and throughput. Novel technological developments have also broadened the scope of measurable molecules beyond RNA-only to many tagged molecules, including surface proteins. With every single experiment potentially generating thousands to millions of cell transcriptomes, across diverse cell types, subtypes, transition phases, or perturbed states, increasing effort has been applied to compiling and documenting the vast amount of data for the scientific community. Thus far, availability of data is generally well enforced, however issues still remain that hinder their reanalysis and reuse.

Single cell data analysis has become increasingly user-friendly and requires minimal investment to successfully conduct. However, much of the analysis time is devoted to annotating the cell types and states identified from unsupervised clustering methods. Recently, numerous tools have been developed to simplify cell type annotation by comparing new single cell datasets to existing reference single cell datasets (Abdelaal et al., 2019; Fu et al., 2020). Additionally, scRNA-seq batch-correction methods, such as Seurat’s integration method, fastMNN, Harmony, and others, enable direct comparison between scRNA-seq datasets at the individual cell level, permitting fine-grained reanalysis and comparison to published datasets (Tran et al., 2020). Reanalysis of single cell datasets however requires cell-type or cell-state annotations to provide interpretable comparisons between the query datasets and reference publication data. Another popular reanalysis method uses marker gene lists or gene signatures for each cell type to generate gene-set module scores for each cell. Gene lists are often incompletely presented but can be easily regenerated if cell-level annotations are available (Aibar et al., 2017; Tirosh et al., 2016).

The Gene Expression Omnibus (GEO) database is one of the largest public repositories for array and sequencing data, providing guidelines and infrastructure aimed at facilitating data archiving and query/review. GEO submissions must include raw sequencing data and processed data files generated by the analysis. However, instructions for single cell sequencing deposition have lagged behind the explosion of technological advances. As a consequence, cell-level annotations are frequently missing from data submissions, severely limiting the accessibility and reproducibility of single cell sequencing studies.

The problem of reproducibility and poor dataset documentation is not unique to single cell datasets and it is a continual challenge to ensure that as new data formats arise proper standards are adopted to make the data accessible and reusable. However, single cell data differs from traditional sequencing data in a manner that challenges existing paradigms for dataset documentation. Data repositories were built at a time when 1 sequencing library generally encompassed only 1 sample. In contrast, single cell studies using popular platforms such as the 10x Genomics chromium or Drop-Seq generate thousands of cells per sample, which each require cell-level annotations for reuse by others. As a consequence, the current dataset deposition standards and guidelines generally do not encourage inclusion of cell-level data necessary to reproduce key conclusions from the studies.

Here we examined public GEO records in an effort to curate databases of cell type gene signatures. We find that a minority (< 25%) of records contain sufficient metadata for replication of published findings. Based on these analyses we suggest that, in addition to count matrices, a metadata table containing relevant cell-level information (e.g., sample, clustering, cell-type, pseudo-time information) should be a required component of single-cell RNA-seq submissions. These metadata tables are integral components of commonly used single-cell RNA-seq analysis workflows (Seurat, SingleCellExperiment, scanpy) and are the basis for many of the figures presented in publications.

## Results

### Public single cell RNA-seq datasets frequently contain insufficient metadata for reanalysis

In an effort to curate reference atlases of diverse cell types, we attempted to download cell-by-gene UMI count matrices and associated cell-level metadata from single cell studies in public data repositories. We found that many studies failed to provide cell-level annotations for the deposited data. To determine how frequently studies contain cell-level annotations, we queried the Gene Expression Omnibus (GEO), which is the most commonly used data repository for single cell studies (used by 78.1% of studies with public data in a curated database of single cell studies) (Svensson et al., 2019).

The only scRNA-seq-specific requirement noted in the current GEO guidelines are for raw data deposition (https://www.ncbi.nlm.nih.gov/geo/info/seq.html). The current GEO requirements for supplemental processed data files however are vague and the guidelines do not reference commonly generated single cell data files. To satisfy the processed data requirement, many studies only provide a cell-by-gene UMI count matrix, which is generated from pipelines such as Cellranger, Kallisto-Bustools, or Alevin (Melsted et al., 2019; Srivastava et al., 2019; Zheng et al., 2017). These submissions do not include important results from the downstream, time consuming analyses performed by the experimental expert, including cluster assignments, UMAP/tSNE projection coordinates, and metadata from cell type classification procedures.

In bulk sequencing data, each sample is the subject of analysis and is described by sample level-metadata. This is, however, not the case for single cell sequencing. A scRNA-seq count matrix is simply an intermediate file format, and does not satisfy GEO’s own stance on processed data, “*defined as the data on which the conclusions in the related manuscript are based*”. The scRNA-seq count matrix is comparable to BAM/SAM/BED formats in bulk RNA-seq experiments, which also do not enable direct reproduction of conclusions and are not considered acceptable processed data by GEO. The lack of single cell-specific guidelines leads to reproducibility issues, as studies that only provide count matrices require significant domain-specific expertise to associate per-cell RNA counts with the cell-types or phenotypes described in or central to the associated publication.

GEO records derived from bulk sequencing methods have queryable standardized terms (e.g. RNA-seq, ChIP-Seq, FAIRE-seq). However, to assess the extent of missing cell-level annotations, we crafted an optimized query string to recover single-cell experiments from GEO because there is no specific annotation associated with single cell datasets. A query string of *“expression profiling by high throughput sequencing” AND (“single nuclei” OR “single cell” OR “scRNAseq” OR “scRNA-seq” OR “snRNAseq” OR “snRNA-seq”)* coupled with further keyword filtering using the GEOquery R package returned 3,476 GEO entries (after merging superseries). These included 97.5% of the GEO studies previously manually curated in Svensson et al, supporting the validity of our query method (Svensson et al., 2019). We then programmatically identified supplemental files with names containing the strings “meta, “annot”, “type”, and “clustering”, as well as R and python data formats “rds”, “rda”, “rdata”, “loom”, and “h5ad”. Only 12.6% of GEO entries contain cell-level metadata (18.0% for entries within the Svensson et al curated database).

To confirm the accuracy of our classification approach we performed manual inspection of randomly selected studies that we identified as single-cell datasets (Table S1). 9.8% of studies that we classified as single-cell datasets were instead other sequencing modalities (e.g. bulk RNA-seq), highlighting the importance of having standardized metadata terms to identify single-cell sequencing datasets. Of the remaining true single-cell studies, 6.4% (10 / 156) contained metadata files that were missed by our automated classification. Based on this analysis we estimate that at most 25% of studies deposited in GEO contain cell-level annotations. GEO records do not have single-cell specific library preparation metadata terms (e.g. Smart-Seq2, Drop-Seq, Fluidigm-C1) which limited our ability to programmatically identify studies that deposited each cell as an independent record. These studies may have included relevant cell-level annotations, however the absence of a standardized metadata term (e.g. cell-type) prevented systematic examination of the annotations in these records.

Further exploring the GEO entries with publication information linked through GEO and PubMed, we found that the percentage of metadata-containing entries have slowly improved with time, as pipeline standards matured and awareness of this issue has grown. However, even for studies published in 2020, the fraction with metadata remains at a frustratingly low 20.8% (**Fig. 1A**). In addition, the issue is widespread through various journals of every family and tier (**Fig. 1B**). While enforcement of data deposition through journals has been highly effective at improving data accessibility, once again the lack of specific guidelines towards scRNA-seq supporting information hurts the overall goal.

**Figure 1.**
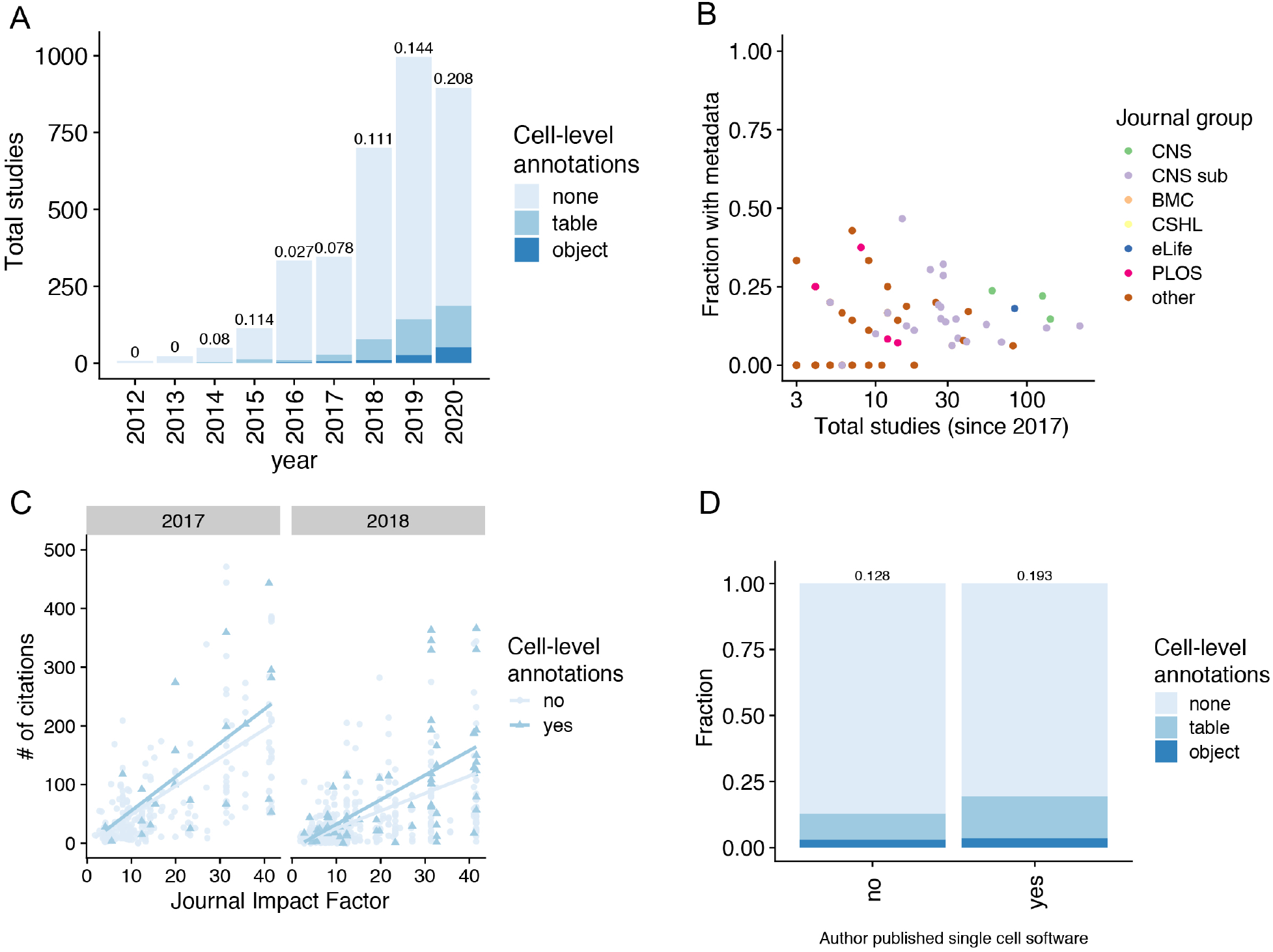
The majority of single cell sequencing datasets do not have cell annotations. (A) Number of single cell datasets in GEO annotated with the proportion that contain cell-level metadata per year, either as plain text tables or binary objects. (B) Fraction of studies published in each group of journals compared to the total number of studies published by each group. (C) Comparison of the number of citations for studies containing or lacking cell-level metadata in 2017 or 2018. (D) Fraction of studies, since 2017, containing cell-level annotations published by authors with a previous publication of a single cell-related software tool.

Next, to corroborate with our own data analysis experience, we explored whether publications with annotated per-cell metadata potentially lead to more citations by facilitating minimal-effort comparison of reported data and cell type gene-expression signatures to new experiments (**Fig. 1C**). Without rigorous statistical testing, due to the limited number of metadata-containing studies and the numerous confounding factors affecting citations, we note that we observe a general trend encouraging the habit of presenting metadata. We also examined datasets deposited by authors on publications describing scRNA-seq informatics classification tools. The tools developed by these authors generally require cell-level metadata, and therefore we hypothesized that submissions by developers of classification tools would be more likely to include cell-level metadata. We identified these authors by querying software curated by the scRNA-tools database, and discovered that GEO entries associated with these authors, who likely most appreciate the value of cell metadata, tend to have better, yet still limited, cell metadata deposition (**Fig. 1D**).

### Metadata is already incorporated in major analysis platforms and is trivial to include with GEO submissions

Cell level annotations are a critical component of single cell RNA-seq analysis. As analysis tool suites have matured, they have converged on similar data storage and annotation solutions. Current common single cell RNA-seq analysis data structures all incorporate cell-level annotations in a table-like format, either as an R data.frame or a pandas DataFrame in Python (**Fig. 2A**). These standard data tables are populated with annotations during the course of the analysis and are trivial to export from the analysis suites (**Fig. 2B**). The coordinates of dimensionality reductions (UMAP, tSNE) can also be easily appended to these tables. Including these coordinates allows users to replicate these projections which are frequently the most common visualization in publications using single cell RNA-seq, but are also not likely to be reproduced upon reanalysis due to the non-deterministic nature of the algorithms.

**Figure 2.**
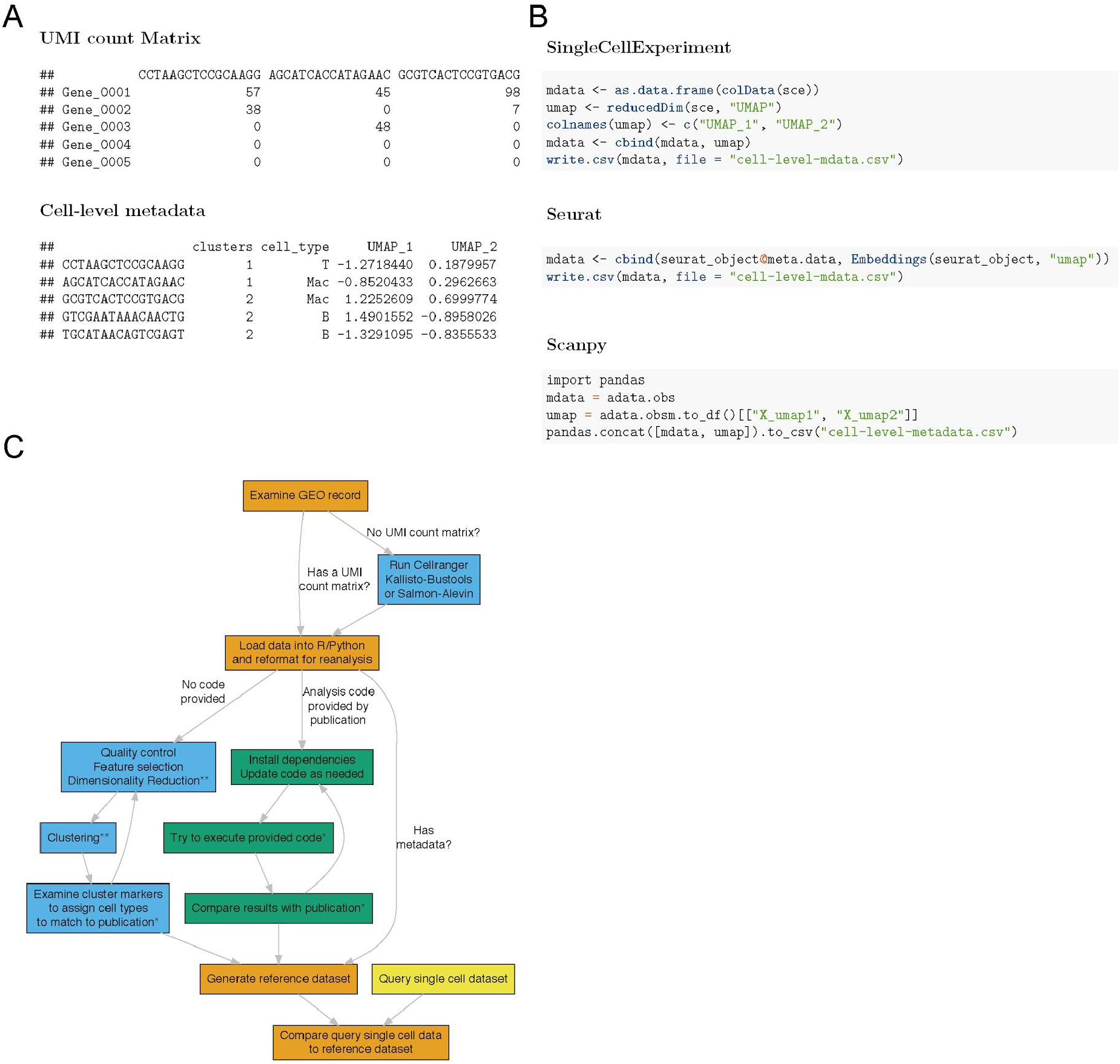
Studies that do not report cell-level metadata are difficult to reproduce. (A) Suggested processed data files for inclusion with single cell studies. (B) Example code for exporting cell-level annotations from popular single cell analysis platforms. (C) Workflow for regenerating cell type or gene-expression signatures from public datasets. * indicates a step requiring an analyst to make subjective decisions ** indicates a step that includes a non-deterministic algorithm.

### Single cell RNA-seq studies cannot be replicated without metadata

In the absence of per-cell metadata, the effort, time, and field-specific expertise required to compare cell subpopulations described in a publication to new single-cell datasets is dramatically increased. Instead of easily leveraging peer-reviewed expertise contained in the cell-level metadata, researchers are forced to rerun pipelines, which can take several hours, and scour the original text for a handful of gene markers described to assign cell type/state subjectively.

Even with careful reanalysis, several factors can limit the original study’s reproducibility if data has to pass through the entire analysis pipeline (**Fig. 2C**). First, exact parameters are often not reported in the manuscript. Even subtle differences can lead to different downstream results. Second, multiple steps in the analysis pipeline are non-deterministic, including the results from clustering and dimensionality reduction. Finally, rapid development in the scRNA-seq software field leads to rapid deprecation of outdated functionalities and possibly silent yet impactful alterations to the underlying algorithms.

In contrast, datasets that provide cell-level annotations enable rapid reproduction and reuse of key findings. For example, GEO accession GSE137710 contains a metadata file for each sample (e.g. GSE137710_human_melanoma_cell_metadata_9315×14.tsv.gz) with cell-level annotations (e.g. “b_cell”, “melanoma”, “myeloid”, “T/NK”) (Brown et al., 2019). Each cell barcode is properly annotated with the cell-type described in the study, which enables very rapid (< 5 minutes) downstream analyses to compare expression patterns and markers for these newly described cell-types to other single cell datasets.

### Suggestions for improving reliability of public scRNA-seq datasets

Moving forward, we suggest the following remedies to this issue:

### For investigators and reviewers

1. Require that analysts provide a metadata table containing cell-level annotations and, if applicable, a binary object saved from the analysis framework (.rds for R or .h5ad for Python).
2. When reviewing single-cell sequencing studies, ensure that the authors have deposited the proper cell-level metadata alongside the raw data into a suitable repository (e.g. GEO, ArrayExpress)

### For journals

1. Include language about requirements/recommendation for external datasets to contain proper cell-level metadata.
2. Ask reviewers to review material deposited to external data repositories.

### For data repositories

1. Public repositories should introduce a standardized annotation that specifies that the dataset contains single cell data. For GEO, commonly used single cell sequencing methods could be added to the library strategy annotation (e.g. scRNA-seq, snRNA-seq, CITE-seq, etc.).
2. Updating submission guidelines to require metadata with cell-level annotations for single cell dataset submissions. For GEO, this would be accomplished by updating the “Processed data files” requirements to outline required data types for single-cell sequencing submissions (**Fig. 2A**). “*For single-cell sequencing data, in addition to standard count matrices (genes-by-cells), we expect users to deposit metadata with cell-level annotations generated during the course of analysis.”*
3. Encouraging previous depositors of single-cell sequencing data to update their records with cell-level metadata, if it was not included in the original submission.

## Discussion

scRNA-seq, though greatly insightful, is a very recent development and lacks most of the infrastructure support of other high-throughput techniques such as bulk RNA-seq or microarray. This manuscript is not intended to provide the single-cell sequencing equivalent of microarray standards (e.g. MINME, MINSEQE) (Edgar and Barrett, 2006), which has been recently explored with great care and detail (Füllgrabe et al., 2020), but simply aimed to highlight the troubling issue, encourage adoption of reproducible data-deposition practices, and to promote discussion of best practices within the community. Large scale efforts to curate cell atlases are currently underway in the Human Cell Atlas, Allen Brain institute, and Fly Cell Atlas and we hope that the standards implemented in these consortia can provide guidance of best practices for documenting single cell datasets (Füllgrabe et al., 2020).

The number of single cell sequencing studies has exploded in recent years, providing a wealth of new information about cell-types and cell-states. We hope that improved standards for public data deposition will encourage large-scale archiving and integration efforts for single-cell datasets akin to the efforts of databases such as Recount2 generated for bulk sequencing methods (Collado-Torres et al., 2017). Databases of single cell studies, such as PanglaoDB, are useful for archiving existing data, but lack incorporation of published cell-types likely due to the inadequacy of existing cell-level metadata records in public datasets (Franzén et al., 2019). Other databases, such as SHOGoiN, which curates data from the Smart-seq and other cell type-labeled 1-cell-per-sample techniques or Cell BLAST which uses manual curation and reannotation, are greatly limited in the number of usable studies (Cao et al., 2020; Mori et al., 2020). Efforts to produce single cell atlas from public datasets will continue to require time-consuming curation in study-by-study manner until the deposition of machine-readable standardized annotation files becomes common practice in the community.

A growing number of researchers are actively promoting reproducibility and data exploration by presenting interactive data browsers or hosting code and metadata files on open-access repositories such as GitHub. However, not all popular cell browser solutions offer metadata export, and external datasets not linked and documented in GEO are difficult to navigate. Hence, a more guided and standardized effort that will facilitate scientific transparency and communication while requiring very little additional work on the part of authors should be well-received.

## Material and Methods

Analysis code is available on GitHub (https://github.com/rnabioco/someta). The repository is set up to automatically monitor the issue of missing cell metadata and build updated reports periodically via GitHub Actions. With each completed automated analysis, the latest version of combined data is available as an Rds object on GitHub.

### GEO query and parsing

GEO snapshot of Nov-20-2020 was obtained via NCBI E-utility calls using a query string of *“expression profiling by high throughput sequencing” AND (“single nuclei” OR “single cell” OR “scRNAseq” OR “scRNA-seq” OR “snRNAseq” OR “snRNA-seq”)*. Series returned by this query were further analyzed with the GEOquery R package, including further filtering of all descriptive fields by keywords listed above, merging subseries from superseries into a single series where applicable, and extraction of supplemental files names (Davis and Meltzer, 2007).

### Programmatic identification of cell metadata files

To determine which GEO entries contain cell annotation metadata, the following assumptions were made: 1. a standalone metadata file should contain “meta”, “annot”, “type”, or “clustering” in its file name; 2. metadata can also be housed in R and python data formats with the extensions of “rds”, “rda”, “rdata”, “loom”, or “h5ad”.

### Additional publication and journal-level analyses

For GEO entries providing linked PubMed IDs, additional publication information was retrieved using R packages easyPubMed and rcrossref (Chamberlain et al., 2016; Fantini, 2019). Cases where the journal name from PubMed is incompatible with rcrossref records were manually fixed before downstream analysis in R. For analysis of scRNA-seq bioinformatic tool authors, scRNA-tools database and R package rbiorxiv were used (Fraser, 2020; Zappia et al., 2018).

